# Emergent collective locomotion in an active polymer model of entangled worm blobs

**DOI:** 10.1101/2021.07.08.451507

**Authors:** Chantal Nguyen, Yasemin Ozkan-Aydin, Harry Tuazon, Daniel I. Goldman, M. Saad Bhamla, Orit Peleg

**Author notes:** These authors have contributed equally to this work (Chantal Nguyen, Yasemin Ozkan-Aydin). These authors have contributed equally to this work (M. Saad Bhamla, Orit Peleg).

## Abstract

Numerous worm and arthropod species form physically-connected aggregations in which interactions among individuals give rise to emergent macroscale dynamics and functionalities that enhance collective survival. In particular, some aquatic worms such as the California blackworm (*Lumbriculus variegatus*) entangle their bodies into dense blobs to shield themselves against external stressors and preserve moisture in dry conditions. Motivated by recent experiments revealing emergent locomotion in blackworm blobs, we investigate the collective worm dynamics by modeling each worm as a self-propelled Brownian polymer. Though our model is two-dimensional, compared to real three-dimensional worm blobs, we demonstrate how a simulated blob can collectively traverse temperature gradients via the coupling between the active motion and the environment. By performing a systematic parameter sweep over the strength of attractive forces between worms, and the magnitude of their directed self-propulsion, we obtain a rich phase diagram which reveals that effective collective locomotion emerges as a result of finely balancing a tradeoff between these two parameters. Our model brings the physics of active filaments into a new meso- and macroscale context and invites further theoretical investigation into the collective behavior of long, slender, semi-flexible organisms.

## 1 Introduction

Throughout the living world, interactions between individuals, and between individuals and the environment, give rise to emergent collective phenomena across scales: cell migration, flocking birds, schooling fish, and human crowds moving in unison [1, 2, 3, 4]. While most examples of collective behavior occur in regimes without physical contact between individuals, many insect, arthropod, and worm species form dense aggregations, where constituent individuals are in constant physical contact with each other, for the purposes of survival, foraging, migration, and mating [5, 6, 7]. Small-scale interactions between individuals enable emergent functionalities at the group level, such as the formation of adaptive structures, including fire ant rafts [8], army ant bridges [9], and bee clusters [10], that can enhance the survival of the aggregation compared to solitary individuals. These living aggregations, where the constituents exert forces on each other or even entangle their bodies into a single mass [6, 11], are associated with the world of soft active matter, which comprises a wide range of systems in which self-propelled individuals can convert energy from the environment into directed motion [7, 12, 13, 14].

Here, we examine the aggregation and swarming behavior of active polymer-like organisms, such as worms, that are flexible and characterized by their slender bodies (i.e., each possessing a length much longer than the width). Some species of worms can physically braid their bodies into highly entangled aggregations [11, 15, 16, 17]. In this paper, we focus on the *Lumbriculus variegatus*, an aquatic worm also known as the California blackworm, blackworm, or mudworm. Blackworms are approximately 1 mm in diameter and up to 2-4 cm in length and live in shallow, marshy conditions across the Northern Hemisphere [11]. The physiology, neurology, and behavior of individual *L. variegatus* has been extensively studied [11, 18, 19, 20], while its collective behavior has only recently been examined [17]. Blackworms can form entangled, shape-shifting blobs which allow the constituent worms to protect themselves against environmental stressors and to preserve moisture in dry conditions [17]. Recent experiments have quantified the material properties and aggregation dynamics of blackworm blobs, which can contain anywhere from a few to over tens of thousands worms and behave as a non-Newtonian fluid [17].

Most notably, these experiments resulted in the first observation of emergent locomotion in an entangled aggregation of multicellular organisms. While some physically-connected aggregations of worms and arthropods have been observed to demonstrate collective, coordinated movement and migration (e.g., [21, 22]), the blackworm blobs demonstrated collective self-transport in temperature gradients [17]. Under a bright spotlight, the worms remained a single entangled unit as they moved toward cooler environments, but only about 70% of worms moved together as an entangled blob in the absence of the spotlight [17]. It was observed in small blobs that the mechanism of this collective movement lies in a differentiation of activity, with outstretched “puller” worms in the front pulling the coiled, raised “wiggler” worms at the back [17].

Other recent work has investigated the rheology and phase separation in aggregations of a similar organism, *Tubifex tubifex*, also called the sludge worm or sewage worm. These worms also form highly entangled blobs in water to minimize exposure to poisonous dissolved oxygen, though collective locomotion has not been observed [23, 16]. It was shown that modeling the aggregation of small worm blobs into larger blobs as the coalescence of droplets undergoing Brownian motion was insufficient to capture the dynamics observed in the experiments [16]. Namely, the diffusion constant of the blobs, which describes how quickly the blobs explore space, was observed to be independent of their size, rather than scaling as the inverse of the blob radius as would have been expected assuming completely random motion; this discrepancy was attributed to the *active* random motion of worms at the surface of the blob. Moreover, it was asserted that a model of collective worm behavior would likely need to account for the self-propelled tangentially-driven motion of individual worms [23].

Motivated by these experiments and insights on aggregations of blackworms and sludge worms [17, 23, 16], we pursue a theoretical model that captures the collective behavior of aquatic worms by linking together local rules governing interactions between individual worms with the emergent macroscale dynamics of the blob. Worms consume energy in order to propel themselves, and as such we look toward the extensive body of research in modeling active polymers and worm-like filaments, where activity can be implemented in different ways, such as via colored noise or a constant tangential force [24, 25, 26, 27, 28]. In general, application of these models has been geared toward biopolymers and unicellular organisms in the microscopic regime, such as actin filaments, microtubules, cilia and flagella, and swarms of slender bacteria [26, 27, 29, 30, 31]. In this paper, we adapt the physics of active filaments within a macroscale, whole-organism context in order to characterize the collective behavior of worm blobs.

Aggregation and swarming has also been observed in the nematode *C. elegans* [32]. Agent-based modeling was subsequently used to elucidate the behavioral rules governing collective *C. elegans* behavior [32], in which individual nematodes were modeled in polymer-like fashion as nodes connected by springs, with the head node undergoing a persistent random walk and the rest of the body following.

Here, we are primarily interested in tangentially-driven active filaments [25, 27], as their behavior is qualitatively similar to that of worms. Such semi-flexible, tangentially-driven filaments demonstrate a rich diversity of behavior. The bending rigidity, activity, aspect ratio, and density of filaments define phases of flocking, spiraling, clustering, jamming, and nematic laning [27].

Drawing upon these models, we model worms as two-dimensional active Brownian polymers, driven by experimental observations of the behavior of single worms (Fig. 1A), worm blobs (Fig. 1B), and the collective locomotion of worm blobs in temperature gradients (Fig. 1C, [17]). We model each worm as a polymer with a tangential self-propulsion force acting only on a portion of the worm designated as the head end, as this qualitatively reflects our observations of worms being more active at the head (Fig. 1D). After developing this single-worm model, we simulate worm blobs via aggregation of multiple identical worms (Fig. 1E) attracted to each other via a strong Lennard-Jones potential. Though our model is in 2-D, we nevertheless observe emergent collective locomotion of the worm blob in a temperature gradient, which arises when the strength of attraction between worms is balanced by the magnitude of the tangential force (Fig. 1F).

**Figure 1:**
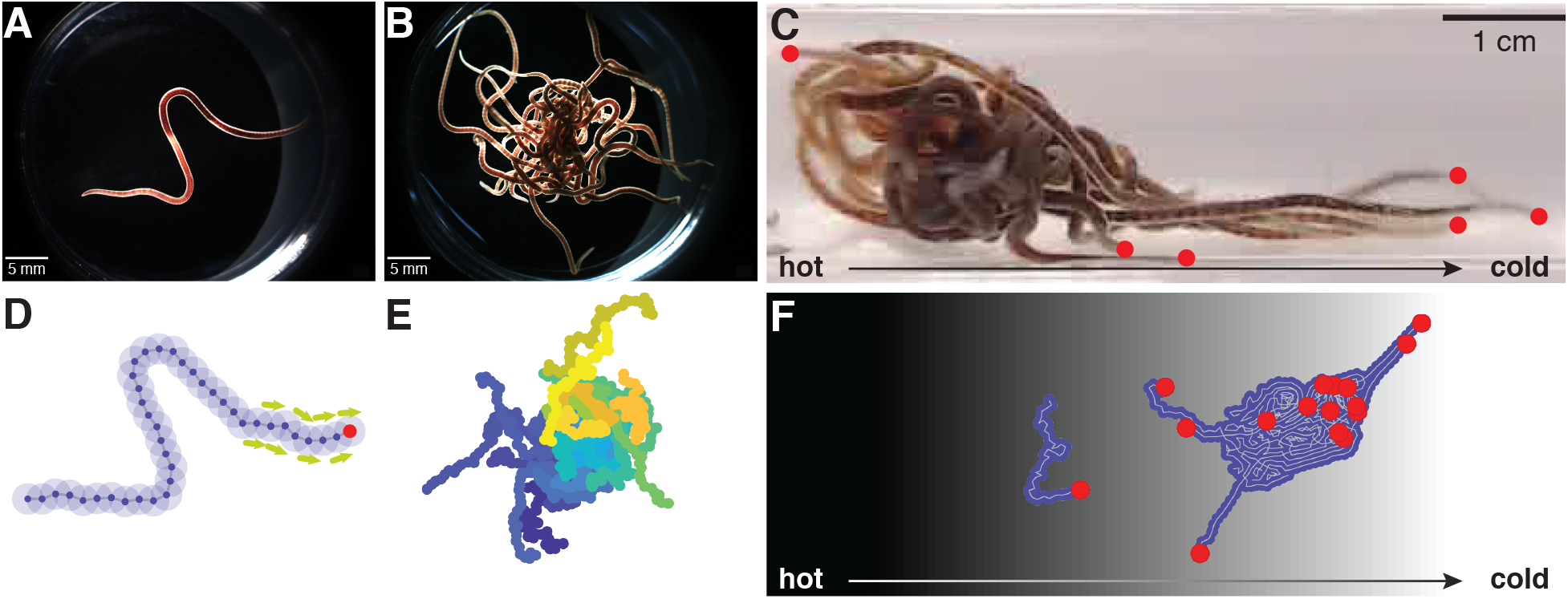
Worm-inspired active polymer model. A: A single California blackworm (*L. variegatus*). Scale bar indicates 5 mm. B: An entangled worm blob consisting of 20 worms. Scale bar indicates 5 mm. C: An entangled worm blob ( 20 worms) in a temperature gradient displays collective locomotion toward the cold side (right), with “puller” worms extending from the front (worm heads marked by red dots). Scale bar indicates 1 cm. D: Polymer model of single worm consisting of 40 monomers connected by springs. The “head” section (with the distal head node indicated by the red dot) of the worm is subject to a constant-magnitude tangential force representing self-propulsion. E: Simulated worm blob consisting of 20 polymers. Each color represents a different worm. F: Simulated worm blob in a temperature gradient with the hotter side on the left (black background) and the colder side on the right (white background) demonstrates collective locomotion toward the cold side. Red dots indicate the head ends of each worm; some worm heads protrude from the bulk of the blob.

## 2 Active polymer model

To construct our model, we begin by modeling a single worm as a polymer: a series of individual monomers linked together by springs of equal length (Fig. 1D). The monomers are subject to three potentials: modified Lennard-Jones (*U*_LJ_), spring (*U*_spring_), and bending (*U*_bending_, described by a harmonic angle potential):

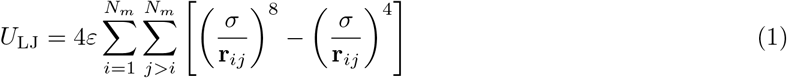

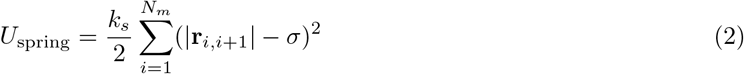

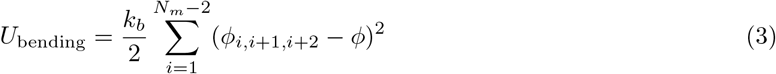

where *N_m_* is the number of monomers per chain, **r**_*ij*_ = **r**_*j*_ − **r**_*i*_ is the vector between the positions of monomers *i* and *j*, *σ* is the equilibrium length of the spring connecting two adjacent monomers, *k_s_* is the spring constant, and *k_b_* is the bending stiffness. The bending potential *U*_bending_ is computed for every consecutive triplet of monomers *i, i* + 1*, i* + 2 whose connecting springs form an angle *φ_i,i_*_+1*,i*+2_ = cos^−1^(**r**_*i*+1*,i*_ · **r**_*i*+1*,i*+2_/|**r**_*i*+1*,i*_||**r**_*i*+1*,i*+2_|). *φ*_0_ is the equilibrium angle of each adjacent pair of springs and is set to *π*.

For two monomers with a separation *r* < *σ*, the Lennard-Jones potential serves as an excluded volume potential to prevent the monomers from occupying the same space. For two monomers with separation *r* > *σ*, the potential is weakly attractive. This results in the polymer forming a more coiled-up conformation. The coiling up is offset partially by the bending potential, which acts to straighten out the polymer.

At each step of the simulation, the force on each monomer is computed:

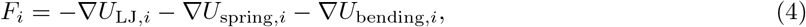

and the position of each monomer is updated via the overdamped equation of motion

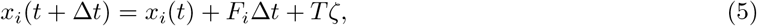

where *ζ* is a variable sampled from the normal distribution 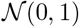 with mean 0 and variance 1, such that *Tζ* represents noise with standard deviation given by a temperature value *T*.

*L. variegatus* cultivated in the laboratory measure approximately 25 ± 10 mm in length with a radius of 0.6 ± 0.1 mm, corresponding to an average length-to-radius ratio of approximately 40. In our model, each pair of monomers is connected by a spring with equilibrium length *σ*, which is also set to be the equilibrium distance at which the Lennard-Jones potential of each monomer has value 0. In our simulations, we model worms that are *N* = 40 monomers long, such that each worm can be considered to have a length of 41 *σ* with radius *σ*, corresponding to a length-to-radius ratio of 41. We also set the spring coefficient *k_s_* = 5000, a relatively high value as worms do not easily stretch along their axis. We also set the bending coefficient *k_b_* = 10, an intermediate value that results in more elongated worms at low temperatures and coiled worms at high temperatures (2A-D). This bending coefficient is also partially offset by a Lennard-Jones coefficient of *ɛ* = 1.

This simulated worm-like polymer is governed by the thermal fluctuations, and as such the polymer exhibits Brownian motion. However, previous studies have indicated that a Brownian depiction does not accurately describe worm behavior [16]. We also observe that simulated worms in this Brownian model demonstrate little exploration of the simulation arena (e.g., low mean squared displacement) at low temperatures, with greater exploration at high temperatures due to large random fluctuations. However, in our observations of real *L. variegatus*, blackworms often demonstrate greater exploration at low temperatures (Fig. 2A). Thus, we add an additional active force that reflects the self-propelled forward peristaltic crawling of the worm (Fig. 1). The force acts with equal magnitude on 8 monomers at one end of the worm, denoted the head end, in the tangential direction determined by averaging the position vectors of the links on either side of the monomer; that is, 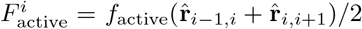. This also reflects our observation that the worms in experiments demonstrate more activity at their heads than from the rest of their bodies.

**Figure 2:**
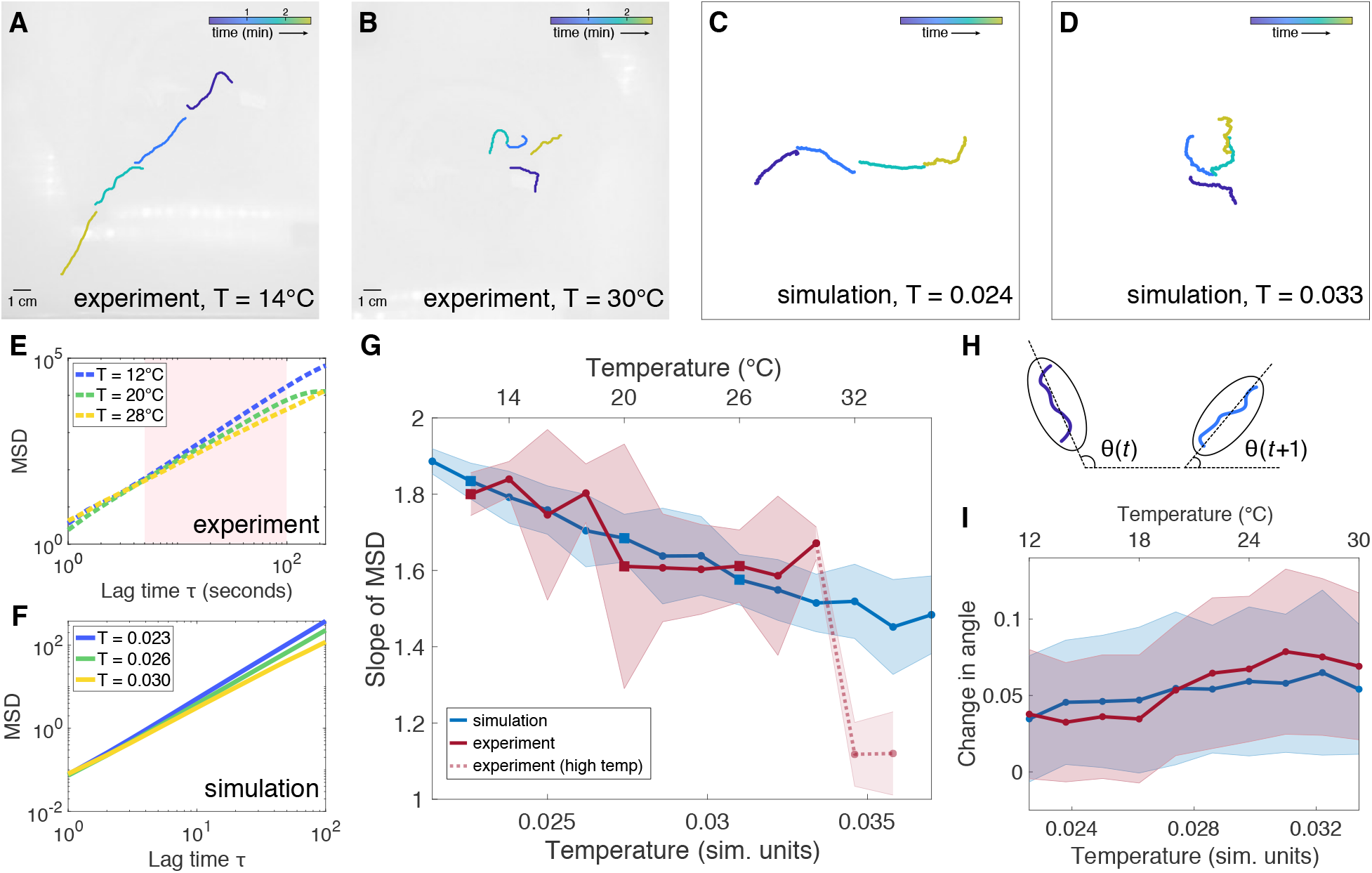
Active polymer model of a single worm. A-B: Snapshots of single worm experiments at *T* = 14°C and *T* = 30°C, respectively. Color bars represent time. C-D: Snapshots of simulated single worm conformations at *T* = 0.024 and *T* = 0.033 (simulation units), respectively. E: Examples of mean squared displacement (MSD) as a function of lag time *τ* from three separate experimental trials at three different temperatures, indicated by the squares in panel G. Because the MSD is not completely linear over the entire range of *τ*, the slope is computed for the region shaded in pink. F: Examples of MSD from three simulations at three different temperatures, indicated by the squares in panel G. G: Comparison of slope SD of mean squared displacement (MSD) as a function of temperature for simulation (blue) and experiment (red, 5 trials). The slope of MSD generally decreases with increasing temperature. Experimental data at *T* = 32-34°C are indicated by a dashed line, as worm physiology is likely to be affected by the high temperature. Squares indicate temperatures at which examples of MSD are plotted in panels E-F. H: The angle *θ* at time *t* is computed by fitting an ellipse to the worm and calculating the angle of the major axis with respect to the horizontal. The average change in angle between consecutive timesteps *θ*(*t* + 1) *θ*(*t*) is used as a measure of worm fluctuations. I: For the trials analyzed in panel G, the average change in angle *θ* increases with temperature for both simulation and experiment.

To fit the parameters of the model to reflect the observed behavior of blackworms, we compare simulations with single-worm experiments (Figs. 2AB). Blackworms were obtained from Aquatic Foods & Blackworm Co. (CA, USA). The worms were cultivated in several boxes (35 × 20 × 12 cm, 25 g of worms per box) filled with spring water (at a height of approximately 2 cm) at ∼4°C for at least three weeks. Worms were habituated to room temperature in a 50 mL beaker with spring water at ∼20°C at least 6 hours prior to experiments. Worms were fed with tropical fish flakes twice a week, and the water was changed one day after feeding them. Studies with *L. variegatus* do not require approval by the institutional animal care committee.

In these experiments, a single worm was placed in the center of a 30 × 30 × 1 cm container filled with water at a height of approximately 0.5 cm. We recorded experiments at water temperatures from 12 to 34°C ± 1°C in increments of 2°C. The worm behavior was recorded at a rate of 2 frames per second for 15 minutes. Video frames were analyzed using MATLAB Image Processing Toolbox (MathWorks, Natick, MA, USA) to extract the position and geometry of the worm. Example trajectories of the tracked worms are animated in SI Movies S1-4 and plotted in SI Figs S1-12.

We observe that at temperatures of 30°C or lower, the worm tends to explore the arena. Often, the worm will travel in a relatively straight path until it reaches the wall of the container, after which it will then continue to explore along the edge of the wall. In some cases, the worm fails to find the wall and continues to explore somewhat erratically. Beyond 30°C, the worm exhibits significantly less exploration, staying close to its original starting position. We attribute this to the temperature being too high for the worm to comfortably explore, and potentially even causing physiological changes to the worm [33]. Above 34°C, the worm is unlikely to survive for more than a few minutes if not seconds.

In our simulations, the self-propelled Brownian polymer remains subject to Gaussian thermal fluctuations. Most noticeably at low *T*, the active tangential force results in the simulated worm moving persistently in a single direction (Fig. 2C). At high *T*, the thermal fluctuations tend to dominate over the bending potential, resulting in a coiled-up conformation of the simulated worm, and as such the individual tangential forces are likely to effectively cancel each other out in direction, resulting in lower overall displacement (Fig. 2D, also see SI Movies S5-7).

To compare simulation and experiment, we examine the mean squared displacement (MSD) as a function of lag time *τ* (Figs. 2E-G and SI Figs. S1-12). Our key observation is that the slope of the MSD, when plotted on a logarithmic scale, differs depending on the temperature. A higher MSD slope indicates that the worm undergoes more directed motion, while a lower MSD slope indicates more diffusive motion, with a slope of 1 representing Brownian motion. In both experiment and simulation, the slope of the MSD generally decreases as temperature increases: at low temperatures, the worm displays near-ballistic movement, which becomes increasingly less directed as temperature increases. Because the worm is confined in the experiments, the MSD is limited by the size of the arena and begins to plateau at large values of *τ*. Hence, we calculate the slope for the regime in which the logarithm of MSD is generally linear, for *τ* between 5 and 100. The simulated worm is not subject to boundary conditions and, at low *T*, will move persistently in the direction set by its initial orientation.

By comparing the slope of the MSD from experiments with simulations, we derive a rough scaling of the temperature between simulation units (*T*_sim_) and degrees Celsius (*T*_exp_): *T*_exp_ = (5000/3)*T*_sim_ − 77/3. In determining this scaling, we excluded experimental data above 30°C, due to the drastic decrease in worm activity at high temperatures. Hence, this scaling is valid only for temperatures between 12 and 30°C inclusive.

While the slope of the MSD captures whether the worm’s motion is directed or random, it does not capture higher-order measures of worm activity. To examine the amount by which a worm fluctuates over time, we calculate the average change in angle of the worm between consecutive timesteps (Figs. 2H-I). The angle *θ* is determined by fitting the smallest ellipse that encloses the worm and calculating the angle of the major axis with respect to the horizontal direction (Fig. 2H). The change in angle increases with temperature, reflecting the greater fluctuations observed in both simulation and experiment.

## 3 Worm blob aggregation

To model a collective system of worms, we retain the dynamics of the single-worm model, but specify a stronger Lennard-Jones potential between monomers of different chains:

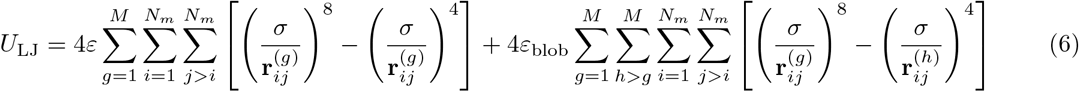

where *M* is the number of worms in the system, *N_m_* is the number of monomers per worm, and the coefficient *ɛ*_blob_ > *ɛ*. We refer to *ɛ*_blob_ as the attraction parameter governing the strength of attractive forces between worms.

We observe in experiments that temperature affects the attachment of worms in a blob, resulting in a transition between a solid-like phase to a fluid-like phase (Fig. 3A). At a low temperature (10°C), a tightly entangled blob remains approximately the same size over the course of several minutes. At a moderate temperature (25°C), the worms spread out slightly, though the blob remains intact; at a high temperature (35°C), the worms quickly disentangle from one another, forming a fluid of detached, coiled worms.

**Figure 3:**
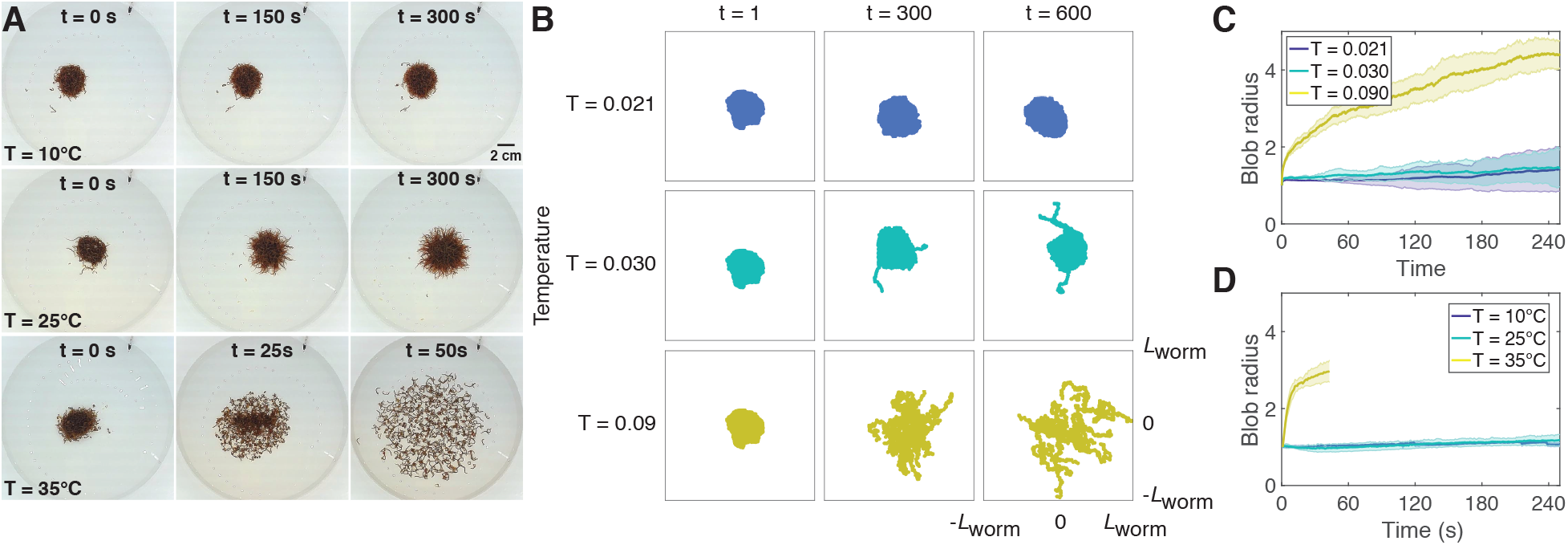
Temperature affects cohesion of worm blobs. A: Snapshots of experimental blobs (*N* ~ 600) at 10, 25, and 35°C. Experiments at *T* = 35°C were performed for only 1 minute since worms begin to die after about 2 minutes at this temperature. B. Snapshots of simulated blobs at different temperatures (row) and time steps (column). Each box is a square with side twice the equilibrium length of one worm (*L*_worm_ = 41*σ*). Each blob contains 20 worms with attachment strength *ɛ*_blob_ = 12. C: Radius ± SD of simulated blob as a function of time for different temperatures (T = 0.021 (blue), 0.030 (green), and 0.090 (yellow)), corresponding roughly to the experimental temperatures used in panel A. For each trial, the radius is normalized to the initial radius at *t* = 0. D: Radius ± SD of experimental blob (*N* ~ 600) as a function of time for different temperatures (T= 10 (blue), 25 (green), 35 (yellow) °C). For each trial, the radius is normalized to the initial radius at *t* = 0.

We simulate worm blobs at three different temperatures, *T* = 0.021, 0.030, and 0.09 (Fig. 3 and SI Movies 8-10). The first two temperatures roughly correspond to 10 and 25°C, respectively, following the temperature scaling described in Section 2. Since temperatures above 30°C result in drastic changes to worm behavior and scale differently than at moderate temperatures, we choose a high simulation temperature of 0.09 to represent 35°C. In these simulations, *ɛ*_blob_ was set to 12, and the active force magnitude was 220.

Each simulation (row of Fig. 3B) begins from the same initial conditions shown in the *t* = 1 column. To generate these conditions, 20 worms were initialized to random positions: the location of the head node was randomly sampled from a square with side length equal to half the worm length, with the angle of the worm sampled from the interval [0, 2*π*). This ensured that the worms were close enough to aggregate into a single blob. The worms were then allowed to aggregate at a low temperature, *T* = 0.02, for a period of 20 time steps. For this preliminary simulation, the Lennard-Jones coefficient *ɛ*_blob_ was set to 20, and the active tangential force was set to zero (i.e., the worms obeyed Brownian dynamics) to facilitate attachment.

At *T* = 0.021, the worms remain in a compact, solid-like blob, demonstrating little activity. At *T* = 0.030, a few worms begin to detach, but most of the worms remain tightly attached. However, at *T* = 0.090, the blob “melts” into a fluid-like state, as the worms separate from each other and disperse across the arena, corroborating the experimental results (Fig. 3A).

## 4 Emergent locomotion and collective thermotaxis

Previous experiments demonstrated the ability of biological worm blobs to undergo emergent collective locomotion in temperature gradients [17]. The blobs exhibited negative thermotaxis, moving from the high temperature side of the gradient to the low temperature side (Fig. 4A). The collective locomotion was enhanced by shining a spotlight on the worms [17]: a worm blob subject to bright light conditions (5500 lux) moved together as an entangled unit, resulting in a higher success rate (over 90%) of worms reaching the cold side. In contrast, worms under low room light conditions (400 lux) did not move as a compact blob, with most disentangling and moving individually, resulting in a lower success rate of thermotaxis (approximately 70%).

**Figure 4:**
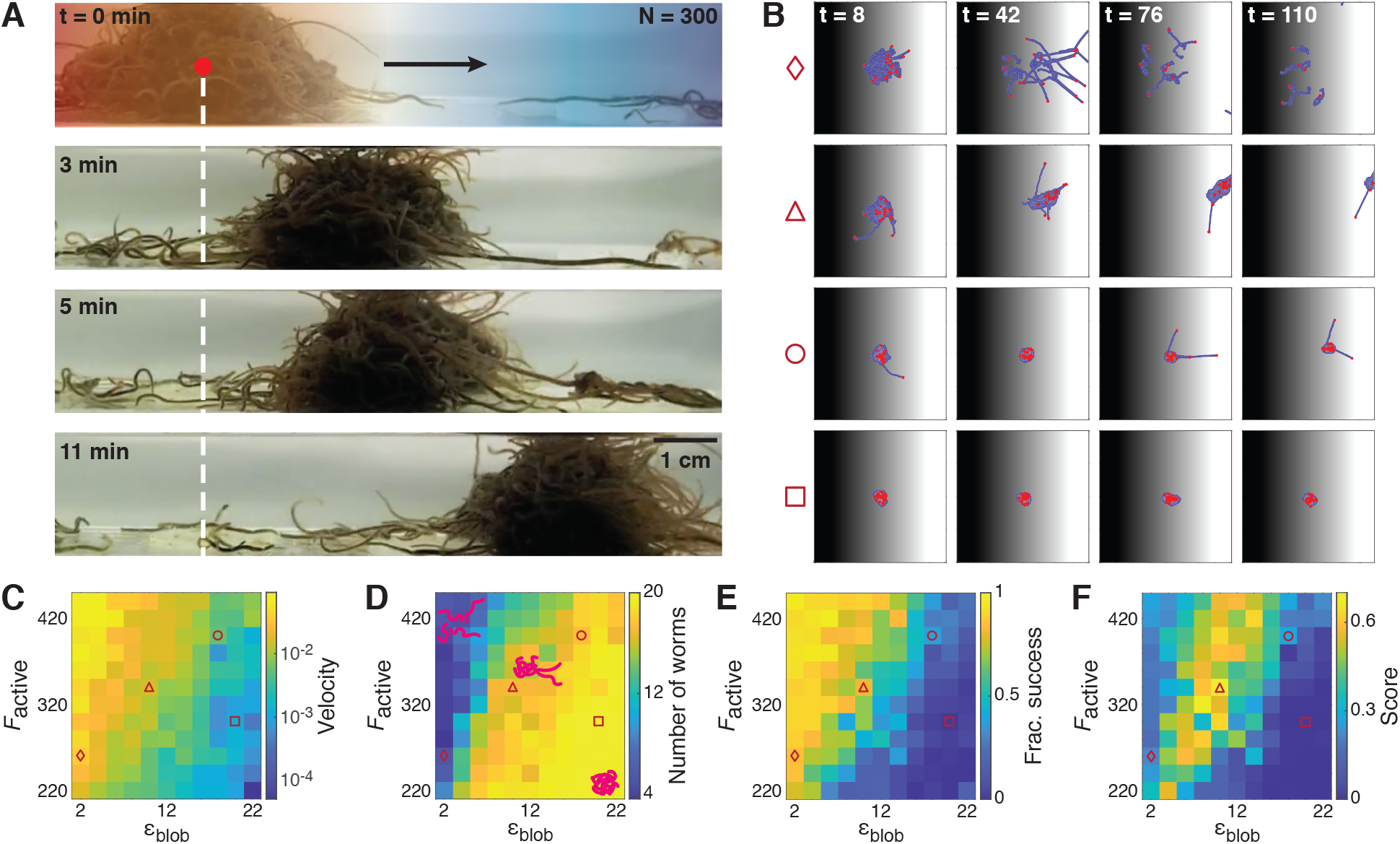
Emergent locomotion in temperature gradients. A: Snapshots of experimental worm blob (*N* = 300 worms) demonstrating emergent locomotion in a temperature gradient (left: high temperature; right: low temperature). B: Snapshots of blob in temperature gradient from *T* = 0.08 (black) on the left to *T* = 0 (white) on the right. Each column corresponds to a different simulation time step (*t* = 8, 42, 76, and 110 respectively). Each row corresponds to different set of attraction parameters *ɛ*_blob_ and active force magnitudes *F*_active_ (diamond: *ɛ*_blob_ = 2, *F*_active_ = 260; triangle: *ɛ*_blob_ = 10, *F*_active_ = 340; circle: *ɛ*_blob_ = 18, *F*_active_ = 400; square: *ɛ*_blob_ = 20, *F*_active_ = 300). Red dots indicate the heads of individual worms. Despite the three-dimensional nature of real worm blobs, our two-dimensional model captures emergent collective locomotion for some combinations of *ɛ*_blob_ and *F*_active_: in the triangle sequence, the majority of worms collectively move from the hot side of the gradient to the cold side, with the heads of some worms extending into the cold side. C: Heatmap of average blob velocity (in worm length per second) for a range of *F*_active_ and *ɛ*_blob_. The velocity is computed for the center of mass of the largest cohesive blob in the simulation. D: Heatmap of the average number of worms (maximum of 20) of the largest cohesive blob for a range of *F*_active_ and *ɛ*_blob_. Cartoons indicate typical configurations of worms for different regimes of parameter space. E: Heatmap of the fraction of successful worms per simulation. Success is indicated by the fraction of worms that reach the *T* = 0 cold side of the gradient by the end of a 1500-timestep simulation. F: Heatmap of the collective locomotion score. The score is given by the product of the blob velocity, blob size, and fraction of success, with each term weighted so that its maximum is 1.

In the single worm case, a simulated worm has no preferred direction in the absence of a gradient; if the temperature is low enough, the worm will move in the tangential direction dictated by its head end, but this direction relies only on the initial orientation of the worm, which is randomly chosen. A temperature gradient will break this symmetry and sets a preferred direction of motion. At high temperatures, the worm’s motion is largely random. However, if, via fluctuations, the head of the worm becomes oriented along the temperature gradient, pointing toward colder temperatures, the tangential forces will cause the worm to move in that direction. The further the worm moves toward lower temperatures, the more it will straighten out, resulting more pronounced ballistic motion.

Here, we simulate blobs in temperature gradients and observe that the level of attachment of worms in a blob depends on a tradeoff between the Lennard-Jones coefficient *ɛ*_blob_ and active force magnitude *F*_active_. The higher the Lennard-Jones coefficient *ɛ*_blob_, the more compact the blob, with worms tightly adhering to one another. Increasing the active force magnitude *F*_active_, on the other hand, increases the likelihood that worms will break apart from the blob.

Simulated blobs with 20 worms were placed in a temperature gradient linearly decreasing from *T* = 0.08 on the left edge of the visualization area to *T* = 0 on the other edge. The visualization area corresponds to a square arena with each edge chosen to be 120 arbitrary units long, approximately 2.5 times the equilibrium length of a single worm. While only this region is visualized, the arena extends indefinitely in each direction, with a constant *T* = 0.08 beyond the left edge and *T* = 0 beyond the right edge.

Each temperature gradient simulation was run from the same initial conditions as described previously in Section 3, where the initial blob aggregated in the absence of a temperature gradient. The same initial blob was used for each simulation and was subsequently placed in the temperature gradient such that its center of mass was located in the center of visualization area, corresponding to *T* = 0.04. We then perform a systematic parameter sweep over the Lennard-Jones coefficient *ɛ*_blob_ between values of 2-22 and the active force magnitude *F*_active_ between 220-420.

In Fig. 4B, we highlight four examples of simulations from different parts of the explored parameter space that illustrate cases in which the blob successfully or unsuccessfully traverses the gradient as a collective. Simulations from a larger sampling of parameter space are also shown in SI Movie S11. If *ɛ*_blob_ is too low (diamond sequence), the worms do not remain attached; if *ɛ*_blob_ is too high (square sequence), the strong attachment forces dominate over the active forces, and the blob remains at its starting position. If *ɛ*_blob_ and *F*_active_ are balanced, this can lead to emergent cohesive locomotion toward the cold side of the gradient (triangle sequence). If *ɛ*_blob_ is slightly larger than *F*_active_, collective locomotion may also occur, but at a slower speed (circle sequence).

In Fig. 4C-E, we compare three quantities as a function of *ɛ*_blob_ and *F*_active_: the velocity of the center of mass of the largest worm blob in the simulation, the size of the largest blob, and the fraction of worms that successfully reach the cold side of the gradient. Generally, we observe that each of these quantities is positively correlated with *F*_active_ and negatively correlated with *ɛ*_blob_, or vice versa.

Fig. 4C is a heatmap of the blob velocity as a function of *F*_active_ and *ɛ*_blob_; as all of the worms in a given simulation may not be attached as a single aggregation, especially for lower values of *ɛ*_blob_, we report here the velocity of the center of mass of the largest cohesive blob as identified using the DBSCAN clustering algorithm [34]. The velocity increases as *F*_active_ increases, but decreases as *ɛ*_blob_ increases.

Meanwhile, Fig. 4D shows a heatmap of the average number of worms in the largest blob, which shows the opposite trend as Fig. 4C: the size of the blob is positively correlated with *ɛ*_blob_ but negatively correlated with *F*_active_. For high *ɛ*_blob_ and low *F*_active_, the blob remains completely cohesive, encompassing all 20 worms. For low *ɛ*_blob_ and high *F*_active_, the worms are less cohesive, with the largest blobs containing down to about 4 worms.

Fig. 4E illustrates the fraction of worms successfully reaching the cold side of the gradient per simulation. This heatmap parallels that of Fig. 4C, showing that the highest proportion of success occurs for low *ɛ*_blob_ and high *F*_active_, and the least successful blobs for high *ɛ*_blob_ and low *F*_active_.

In general, worms are most effective at reaching the cold side when *ɛ*_blob_ is low and *F*_active_ is high. However, they do not move cohesively, with the largest blobs containing between approximately 25-50% of the total worms in the simulation. At the other extreme, when *ɛ*_blob_ is high and *F*_active_ is low, nearly all worms remain in a cohesive aggregation, but the blob demonstrates little to no movement toward the cold side of the gradient, due to the attachment forces dominating over the active motion. We note that for real worms, remaining in a cohesive aggregation is beneficial, especially when there is danger of moisture loss [17]. Moreover, individual blackworms can die within minutes in high temperature environments (above 30°C). Our simulations do not reflect any potential worm death; in some cases, individual simulated worms that have moved toward the hot side of the gradient become “unstuck” via random fluctuations and may eventually find the cold side.

Hence, we seek a regime in which the worms demonstrate a high rate of success at reaching the cold side of the gradient and move relatively quickly while remaining mostly cohesive. To do so, we compute a score for each simulation given by the product of the velocity of the center of mass, largest blob size, and fraction of success, which each of the three terms normalized such that each individual term scales between 0 and 1. All three terms are moreover equally weighted such that the score takes on values between 0 and 1. Fig. 4F illustrates this score as a function of *ɛ*_blob_ and *F*_active_. The tradeoff between *ɛ*_blob_ and *F*_active_ produces a regime in which the highest scores are achieved, along a band that roughly follows the line *F*_active_ = 22*ɛ*_blob_ + 132.

Fig. 5 illustrates a phase diagram corresponding to this function overlaid with example snapshots of blob configurations from corresponding simulations, revealing the rich ensemble of behaviors across the parameter space of *F*_active_ and *ɛ*_blob_. To generate the phase diagram, we fit the score landscape from Fig. 4F to the following function of *F*_active_ and *ɛ*_blob_:

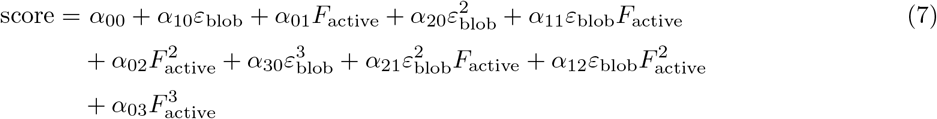

**Figure 5:**
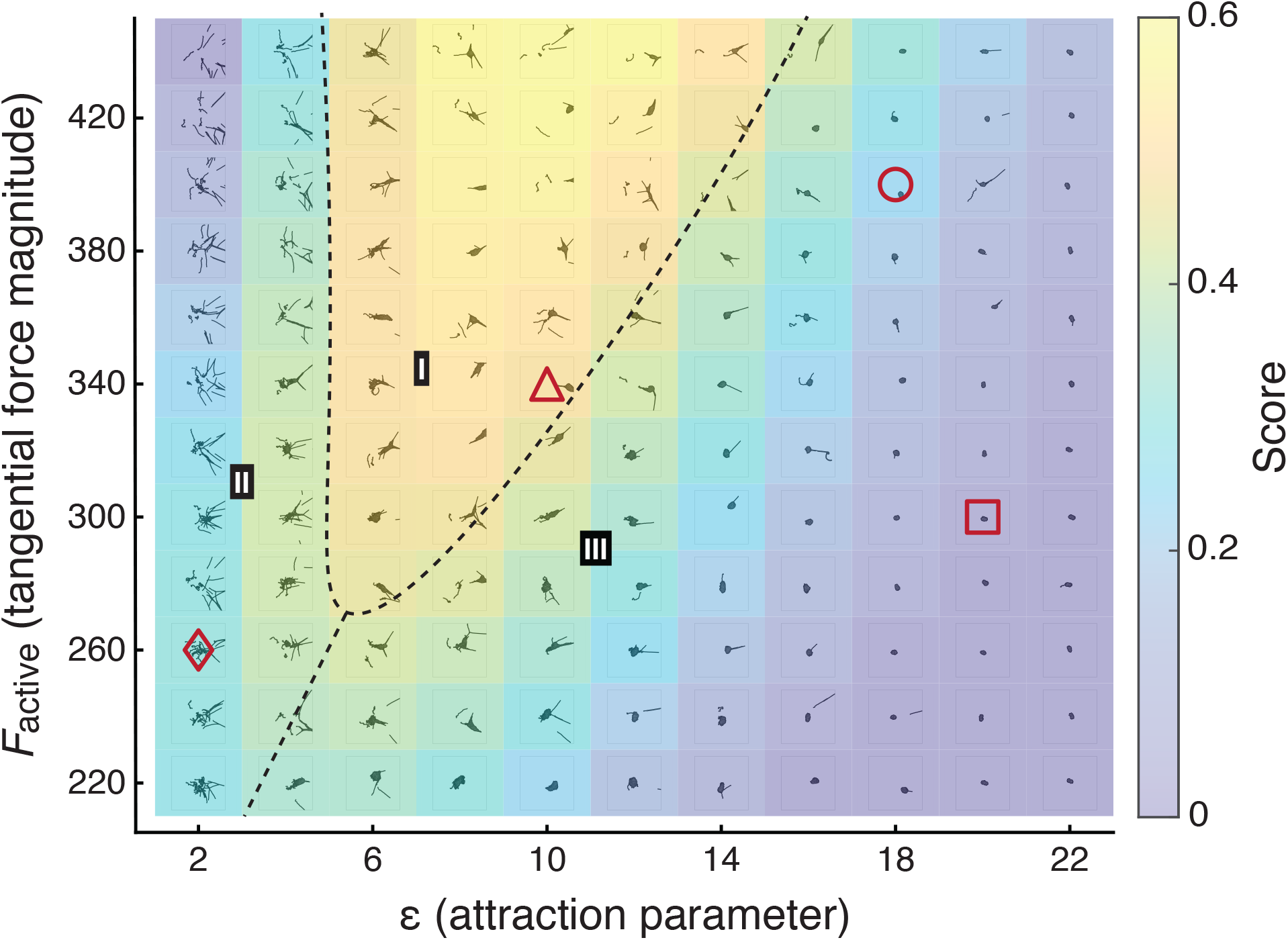
Heatmap of score reveals parameter regime in which the most effective collective locomotion is observed. The score is defined as the product of the blob center of mass velocity, final size of the largest blob, and fraction of worms successfully reaching the cold side of the gradient. The heatmap is divided into three regimes (I-III). I: consistently successful cohesive blob locomotion, reflecting observed emergent locomotion in real worm blobs (Fig. 4A); II: generally unsuccessful blob locomotion, with failure due to dissociation of blobs; III: generally unsuccessful blob locomotion, with failure due to overly strong attachment, resulting in little collective movement. In phases II and III, parameter combinations near the boundary of phase I can intermittently lead to successful collective locomotion. Each subpanel shows an overlaid snapshot at between *t* = 50 and 150 of an example simulation with the corresponding *F*_active_ and *ɛ*_blob_. Red shapes correspond to example sequences shown in Fig. 4B.

The parameters are tabulated in SI Table S1.

The dashed lines in Fig. 5 separate three regions (I-III) characterized by the prevailing collective behavior. In region I, corresponding to the region where the highest scores are achieved, the worms consistently traverse the gradient as a collective. In this region, the emergent collective locomotion reflects what is observed in experiments (Fig. 1C and 4A, [17]). We note that while these real worm blobs inhabit three-dimensional space, our two-dimensional model nevertheless captures collective locomotion. In regions II and III, collective locomotion is generally unsuccessful, though successful cases of collective locomotion are intermittently observed near the boundary with region I. In region II, failures typically occur when the blob dissociates and worms move individually, as *F*_active_ is too high for the corresponding values of *ɛ*_blob_. For failed cases in region III far from the boundary with region I, the worms are too strongly attached and as such the blob does not demonstrate any self-motility and remains near the starting position. Closer to the boundary with region I, the majority of worms may remain attached, with a few worms detaching from the blob and potentially moving toward the cold side on their own. The center of mass of the largest blob either remains close to the origin or drifts slowly to cold side, as here *ɛ*_blob_ is slightly too high compared to *F*_active_.

## 5 Discussion

Following the first observation of collective locomotion in entangled worm blobs [17], we develop a model that places the physics of active, semi-flexible polymers and filaments in the context of the collective behavior of macroscale, multicellular organisms. We model worms as self-propelled Brownian polymers, focusing specifically on the parameter space of aspect ratio, bending rigidity, activity, and temperature that describes the California blackworm, *L. variegatus*, at temperatures between 10 and 35 °C. In the single worm case, the constant-magnitude tangential force *F*_active_ results in a persistent directed motion at low temperatures, with larger fluctuations erasing the persistent motion at high temperatures. In a temperature gradient, this results in a preferred direction of movement from high to low temperatures.

Multiple worms can aggregate into a blob, held together by Lennard-Jones attraction as governed by the attachment strength *ɛ*_blob_. We show that the simulated blob can collectively navigate along a temperature gradient provided that the tangential force and attachment strength are balanced. In a parameter sweep over the attachment strength *ɛ*_blob_ and the magnitude of the tangential force *F*_active_, we observe a tradeoff between the worm velocity and the cohesiveness of the blob. Higher attachment reduces the speed of the blob and hinders collective motion in extreme cases, while a higher force increases the individual worm speed but can result in worms detaching from the blob. We identify the regime where blob movement is “optimal” from a biological perspective – i.e., where the blob quickly moves toward cooler, less dangerous temperatures, while remaining largely cohesive, as worms are less likely to survive on their own outside of the blob – quantified by a score that combines the blob velocity, blob size, and fraction of worms successfully reaching the cold side of the gradient.

We note that a similar tradeoff, between the fraction of successful worms and the blob speed, was illustrated in SI Fig. S12 of [17]. For experiments at four different worm blob sizes (*N* = 10, 20, 40 and 80), it was observed that as the number of worms in a blob increased, the fraction of worms successfully reaching the cold side of the gradient increased as well. However, larger blobs were slower at reaching the cold side. While we have focused on a single blob size in this paper, our model can be expanded to other system sizes and can be used to make experimentally testable predictions of how the number of worms affects collective locomotion. Moreover, future experiments can involve altering the activity of individual worms (e.g. by adding alcohol to the water) and/or their attachment strength (e.g., by manipulating light conditions as observed in [17]) in order to test our predictions on these parameters’ effect on blob motility.

Currently, the parameters of our model are chosen such that there is qualitative agreement between the behavior of the simulated worm and observed *L. variegatus*. However, we expect that our model should be broadly generalizable to describe other long, slender, flexible organisms including annelids and nematodes. Future work will investigate the effect of the aspect ratio of individual worms on their collective behavior in temperature gradients. For instance, *T. tubifex* are a similar length to blackworms but are about a quarter as thick, though collective locomotion in *T. tubifex* has not been observed. It was also observed that terrestrial worms such as common earthworms (*Lumbricus terrestris*) and red wigglers (*Eisenia fetida*) form blobs in air [17], but collective locomotion in such worms remains to be investigated. Moreover, in the limit of an aspect ratio of 1, the polymer picture reduces to that of a single round particle. Such a model can be useful to describe aggregations of organisms that more closely resemble particles rather than filaments, such as ants and bees; future research can work toward a unified model that captures collective behavior along the gradient between active particles and filaments.

A primary limitation of our current model is its two-dimensionality. While we are able to capture collective behavior of active worm-like polymers, in reality, blackworms form blobs that are three-dimensional in nature. In sufficiently deep water, small blackworm blobs are hemispherical (Fig. 4A). Future work will generalize the current model to three dimensions, which will also allow us to explicitly model the physical entanglement of polymers. Entanglement and reptation in polymer melts and solutions has been extensively examined for decades (e.g. by de Gennes [35]). More recently, non-equilibrium polymeric fluids containing active polymers have come under focus, as these systems cannot be explained by statistical-mechanical theories [26, 30, 36]. For instance, Manna and Kumar showed that in a confined volume, contractile active polymers spontaneously entangled, and moreover that this entangled state was stable for any volume fraction of polymers [36]. Meanwhile, for extensile active polymers, they observed a phase transition between disentanglement and entanglement governed by the activity and volume fraction.

By simulating entangled active polymers, we can more closely examine the mechanisms by which blackworm blobs collectively locomote: the differentiation of activity whereby worms at the front are elongated and pull the clump of coiled worms at the back. In particular, we can examine the role of trailing “wiggler” worms that lift themselves off the surface, potentially to reduce friction, which cannot be probed currently with our 2-D model. In experiments, differentiation of activity has only been explicitly observed in small blobs containing on the order of tens of worms, where such differentiation of activity can be seen by eye [17]. These observations were validated by force cantilever experiments, which demonstrated that a few worms were able to exert a force strong enough to pull the blob, and by simple robophysical experiments, in which a blob of entangled small robots could only move as a unit if the group was divided into a few robots that use a “crawl” or “wiggle” gait while the rest remain stationary, as opposed to all crawling or all wiggling [17]. In future simulations, we aim to simulate 3-D entangled worm blobs in order to elucidate whether this collective motion mechanism remains valid as blob size increases.

Here, we have examined the collective dynamics in a general system of active filament-like worms, focusing on a section of parameter space chosen to reflect blackworm behavior. However, real three-dimensional blackworm blobs also exhibit properties that are not captured in our model. For instance, in a surface in air, blackworms form a hemispherical blob to maximize moisture retention; they will also spread out in long “arms” in order to search for moisture and shrink back into a hemisphere if no moisture is found [17]. To more precisely describe this particular biological system, our current model could be expanded to explicitly incorporate rules that describe worms’ sensing of their local environments. Indeed, the interplay between individual sensing and interaction with the environment, coupled with interactions between worms in close proximity, leads to fascinating emergent collective phenomena such as this cooperative searching behavior.

In conclusion, we have developed a model that examines active polymers in the context of entangled living systems much larger than the scale of cytoskeletal, cellular, and other biological systems typically described within similar frameworks. We subsequently identified a regime wherein effective collective locomotion emerges as a result of balancing the tradeoff between directed activity and attachment of individuals. While the experimental observations of the California blackworm in particular have driven our current work, our research opens up avenues for new experiments and theoretical investigations of the collective behavior of long, slender organisms at the meso- and macroscales.

**Table 1:** List of model parameters and corresponding ranges of values used in simulations.

| Parameter | Description | Range of values |
| --- | --- | --- |
| $N_m$ | Number of monomers | 40 |
| $\sigma$ | Equilibrium distance between monomers | 1.189 (arb. units) |
| $\varepsilon$ | Lennard-Jones coefficient, single worm | 1 |
| $k_s$ | Spring constant | 5000 |
| $k_b$ | Bending stiffness | 10 |
| $F_{\text{active}}$ | Self-propulsion force magnitude | 220-440 |
| $\varepsilon_{\text{blob}}$ | Lennard-Jones coefficient for blob (attraction parameter) | 2-22 |

## Acknowledgements

D.I.G. acknowledges funding support from ARO MURI award (W911NF-19-1-023) and NSF Physics of Living Systems Grant (PHY-1205878). M.S.B. acknowledges funding support from NSF Grants CAREER 1941933 and 1817334.

