## Supplementary Material for "Emergent collective locomotion in an active polymer model of entangled worm blobs"

July 8, 2021

### 1 Descriptions of SI Movies

**SI Movie S1.** Movie of tracked worms from three separate experimental trials, at 16, 22, and 30°C, respectively, overlaid onto the same figure. Movie is sped up 8x from actual data.
**SI Movie S2.** Tracked worms from five experimental trials, all at 16 °C, overlaid onto the same figure. Movie is sped up 8x from actual data.
**SI Movie S3.** Tracked worms from five experimental trials, all at 22 °C, overlaid onto the same figure. Movie is sped up 8x from actual data.
**SI Movie S4.** Tracked worms from five experimental trials, all at 30 °C, overlaid onto the same figure. Movie is sped up 8x from actual data.
**SI Movie S5.** Side-by-side comparison of experimental worm at 16°C and simulated worm at corresponding $T = 0.025$.
**SI Movie S6.** Side-by-side comparison of experimental worm at 22°C and simulated worm at corresponding $T = 0.029$.
**SI Movie S7.** Side-by-side comparison of experimental worm at 30°C and simulated worm at corresponding $T = 0.033$.
**SI Movie S8.** Simulated 20-worm blob at a constant $T = 0.021$.
**SI Movie S9.** Simulated 20-worm blob at a constant $T = 0.030$.
**SI Movie S10.** Simulated 20-worm blob at a constant $T = 0.090$.
**SI Movie S11.** Worm blob simulations from a sample of the parameter space explored in the main paper. Each panel corresponds to a separate simulation of 20 worms starting from the same initial conditions. Background gradient corresponds to a temperature gradient of $T = 0.08$ on the left (black background) to $T = 0$ on the right (white background). Each row corresponds to the same value of tangential force magnitude $F_{\text{active}}$, and each column the same value of the attachment parameter $\varepsilon_{\text{blob}}$.

### 2 Supplementary Tables and Figures

| Coefficient | Value |
|---|---|
| $\alpha_{00}$ | -0.75 |
| $\alpha_{10}$ | 0.0022 |
| $\alpha_{01}$ | 0.0088 |
| $\alpha_{20}$ | -0.0057 |
| $\alpha_{11}$ | 0.00025 |
| $\alpha_{02}$ | $-2.4 \times 10^{-5}$ |
| $\alpha_{30}$ | 0.00029 |
| $\alpha_{21}$ | $-2.3 \times 10^{-5}$ |
| $\alpha_{12}$ | $6.1 \times 10^{-7}$ |
| $\alpha_{03}$ | $1.4 \times 10^{-8}$ |

Table 1: Values of coefficients in Eq. 7 of the main text.

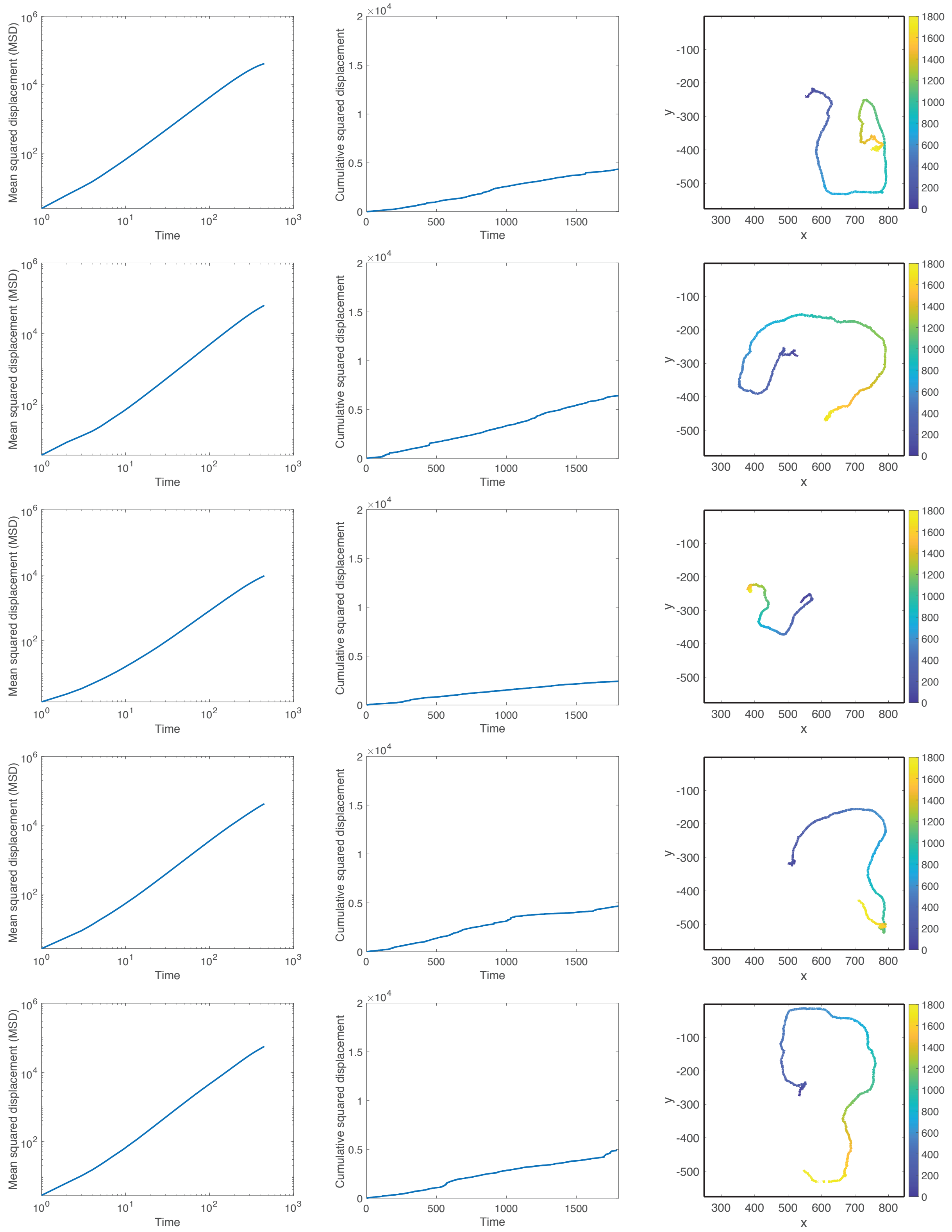


Figure 1: **Single worm experiments at $T = 12°C$.** Left column: mean squared displacement; middle column: cumulative squared displacement; right column: trajectory of worm. Each row represents a different 15-minute trial.

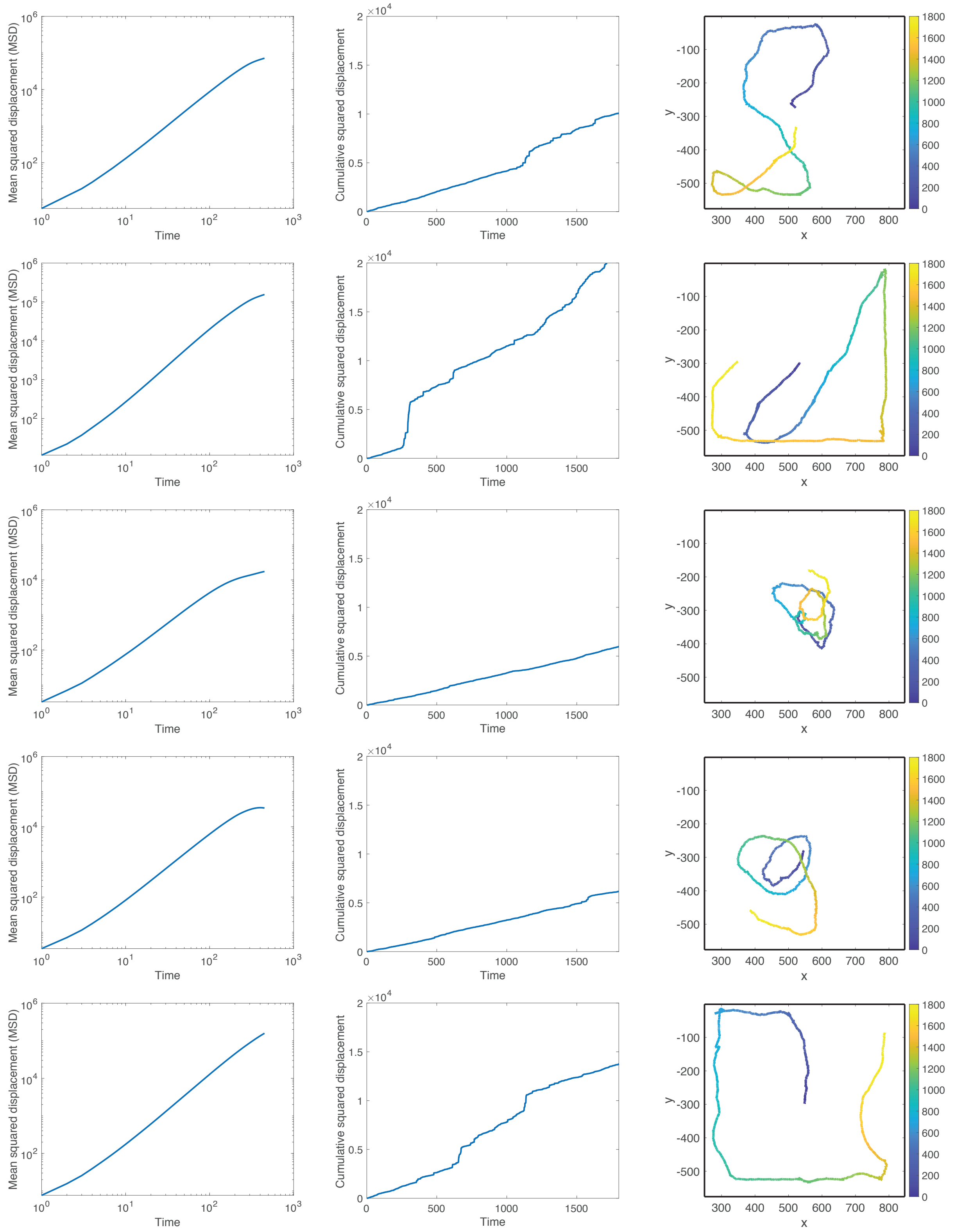


Figure 2: **Single worm experiments at $T = 14°$C.** Left column: mean squared displacement; middle column: cumulative squared displacement; right column: trajectory of worm. Each row represents a different 15-minute trial.

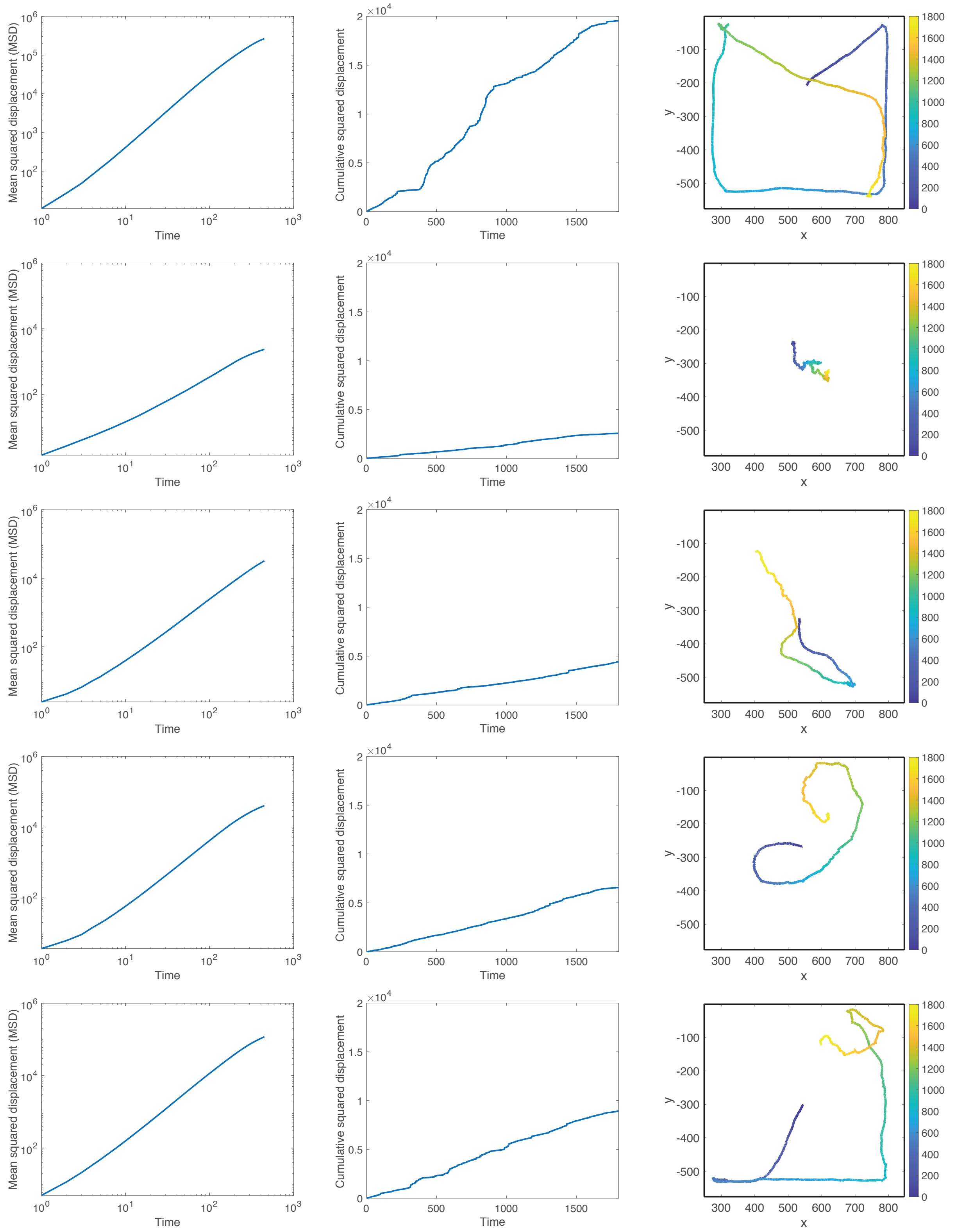


Figure 3: **Single worm experiments at $T = 16°C$.** Left column: mean squared displacement; middle column: cumulative squared displacement; right column: trajectory of worm. Each row represents a different 15-minute trial.

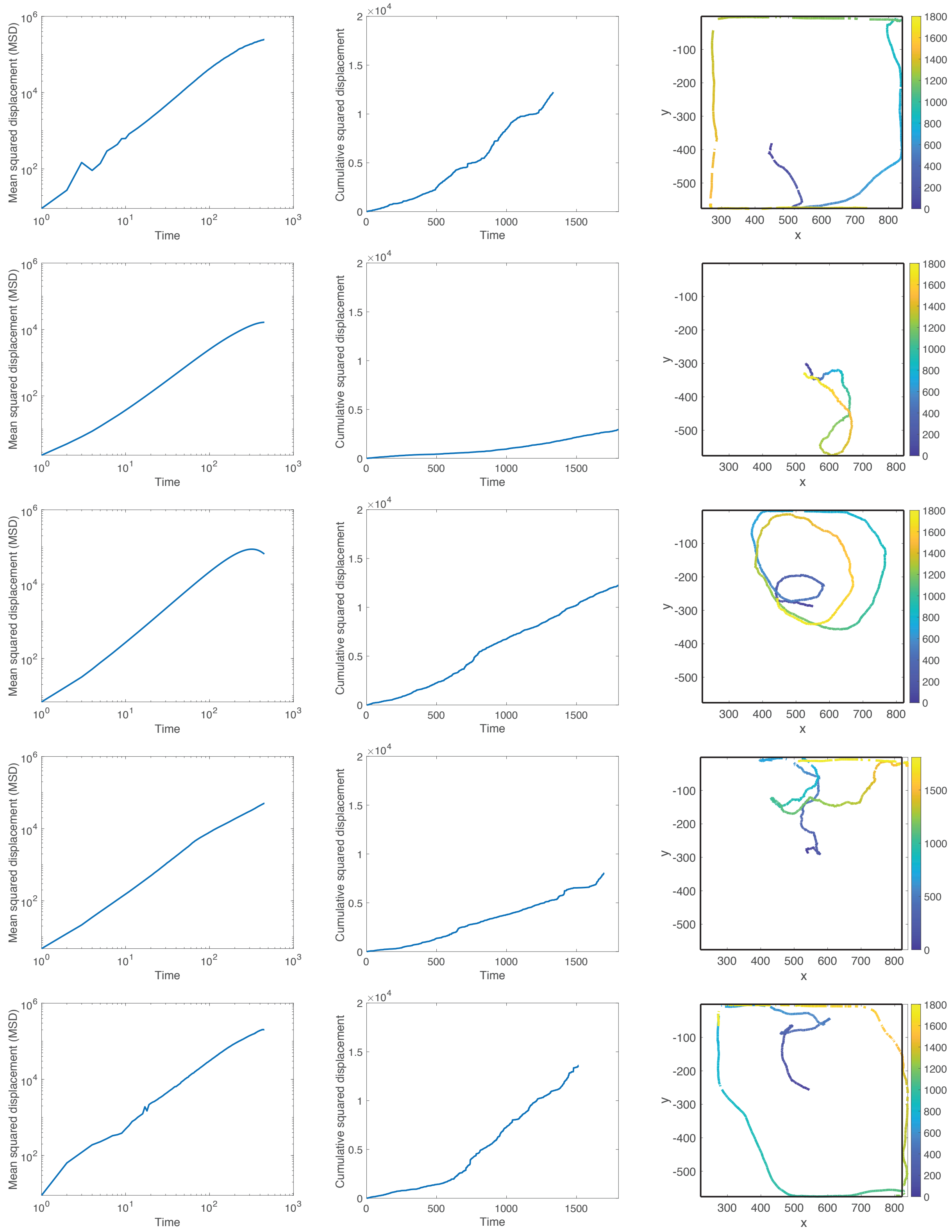

Figure 4: **Single worm experiments at $T = 18°C$.** Left column: mean squared displacement; middle column: cumulative squared displacement; right column: trajectory of worm. Each row represents a different 15-minute trial.

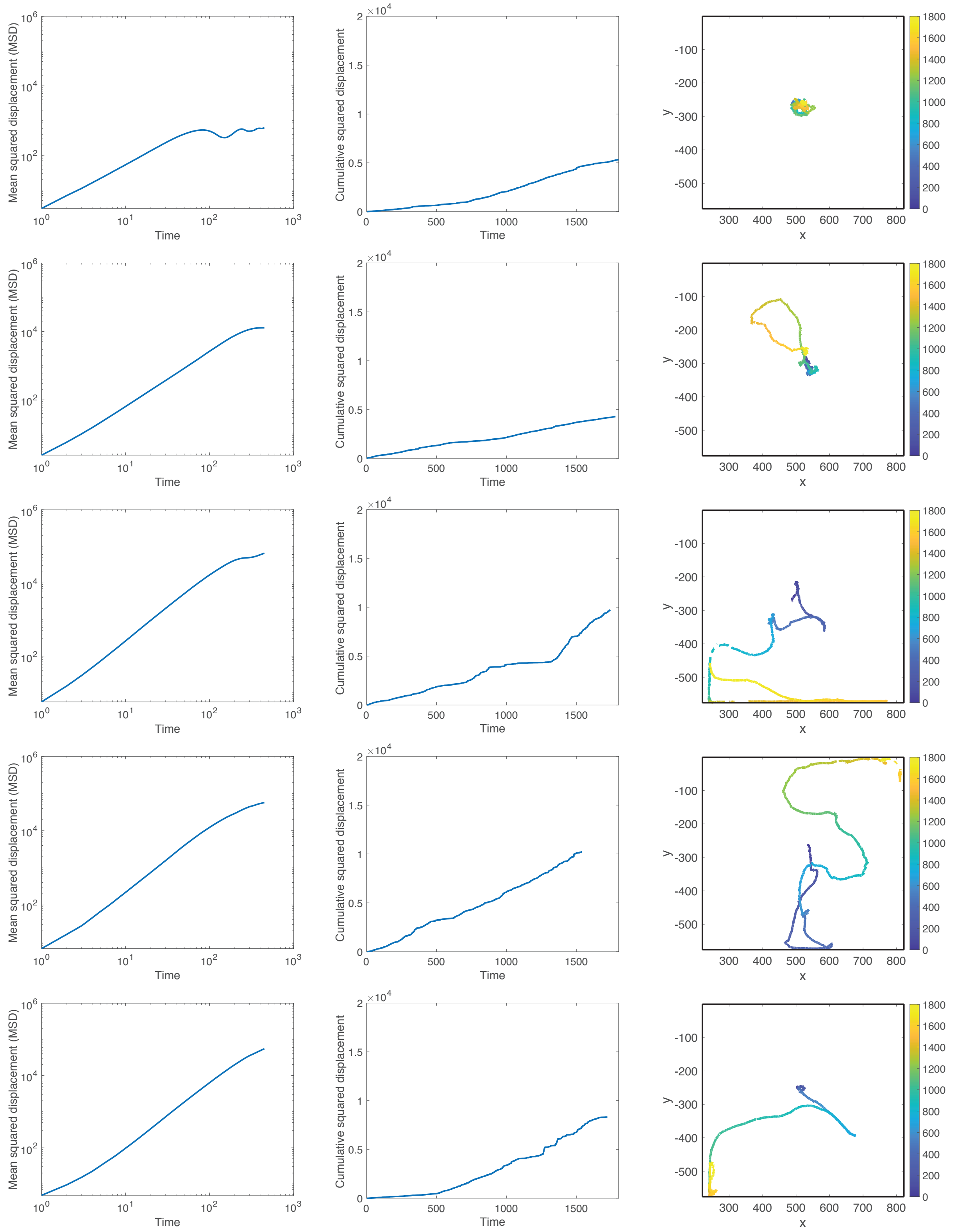


Figure 5: **Single worm experiments at $T = 20°C$.** Left column: mean squared displacement; middle column: cumulative squared displacement; right column: trajectory of worm. Each row represents a different 15-minute trial.

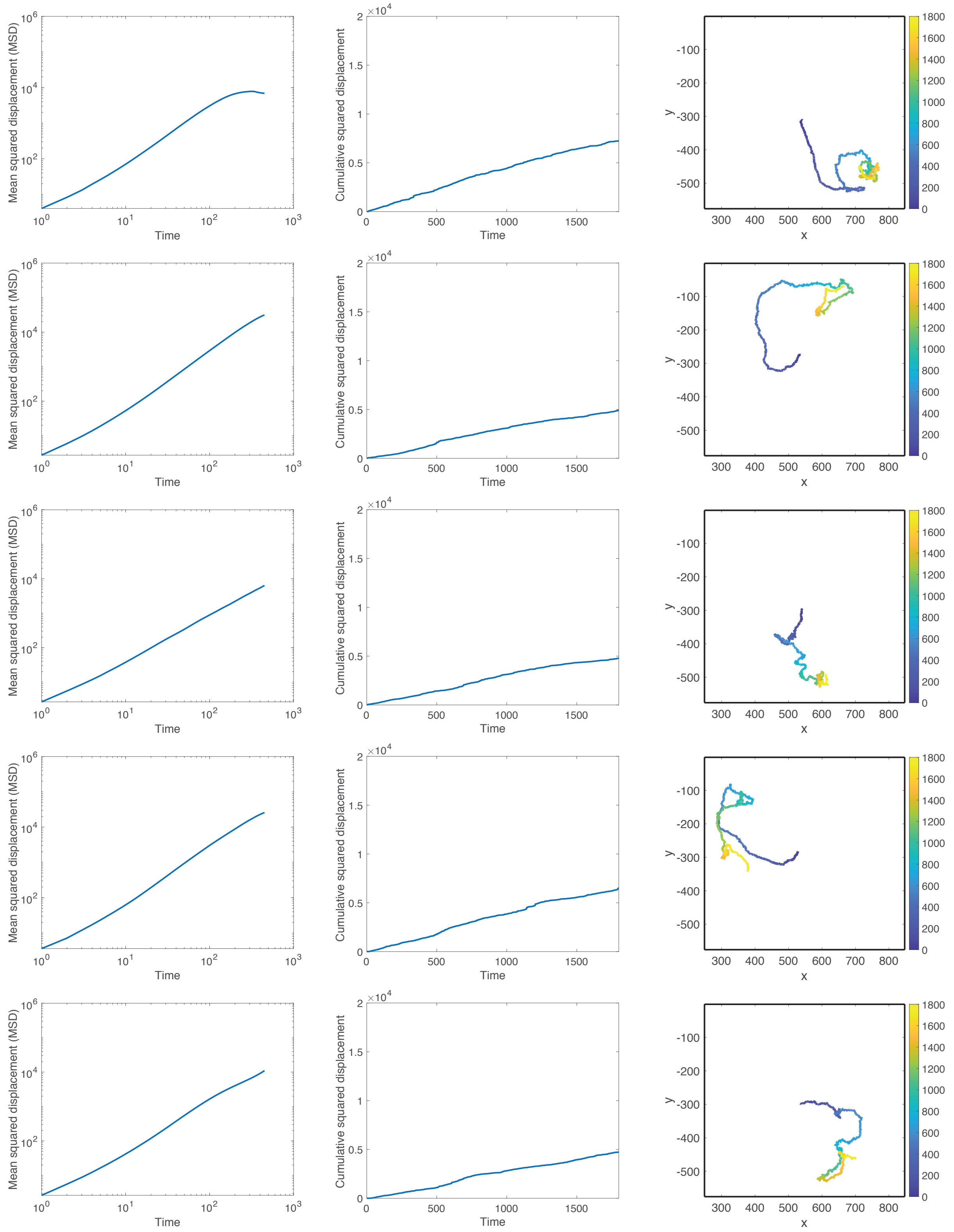


Figure 6: **Single worm experiments at $T = 22°$C.** Left column: mean squared displacement; middle column: cumulative squared displacement; right column: trajectory of worm. Each row represents a different 15-minute trial.

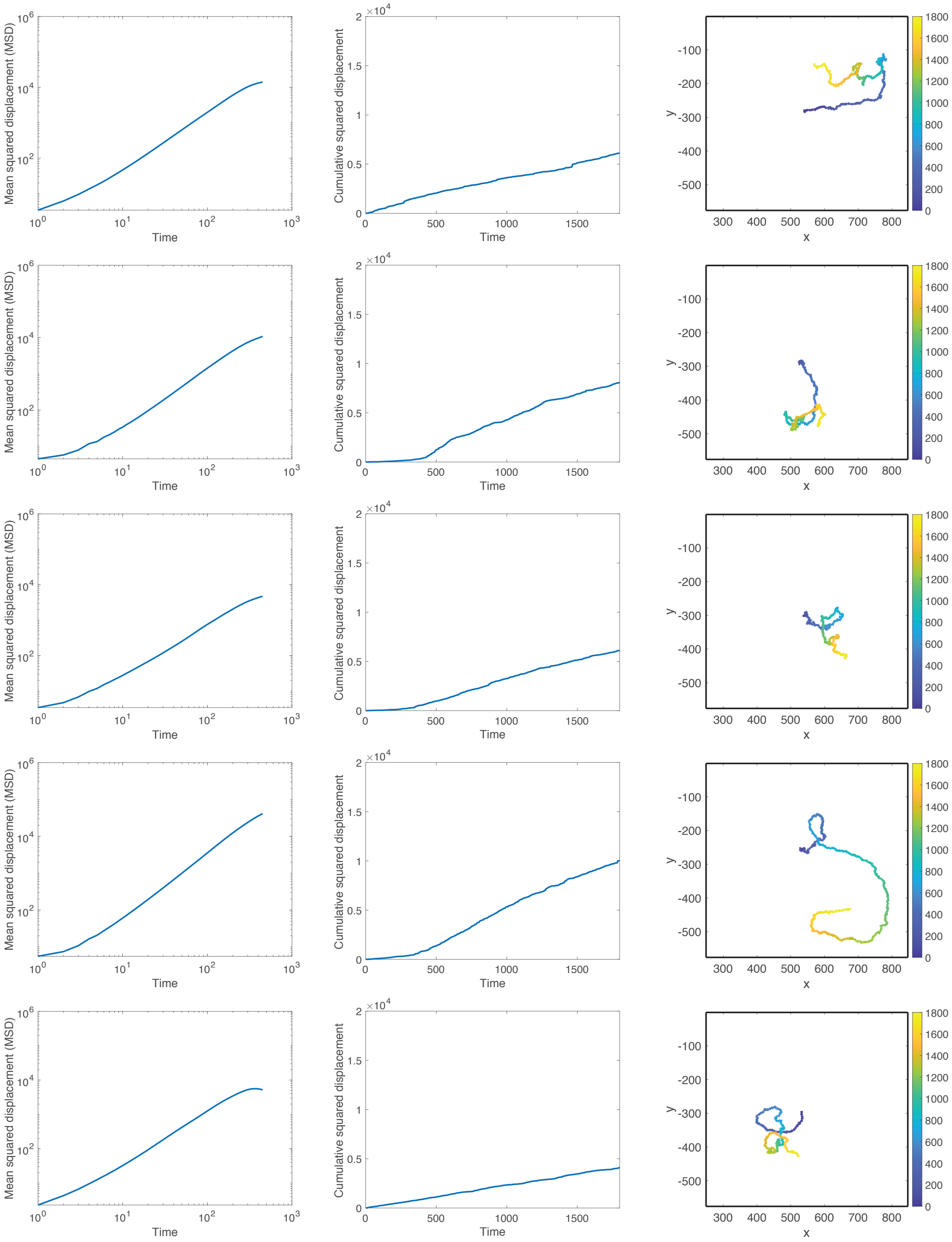


Figure 7: **Single worm experiments at $T = 24°$C.** Left column: mean squared displacement; middle column: cumulative squared displacement; right column: trajectory of worm. Each row represents a different 15-minute trial.

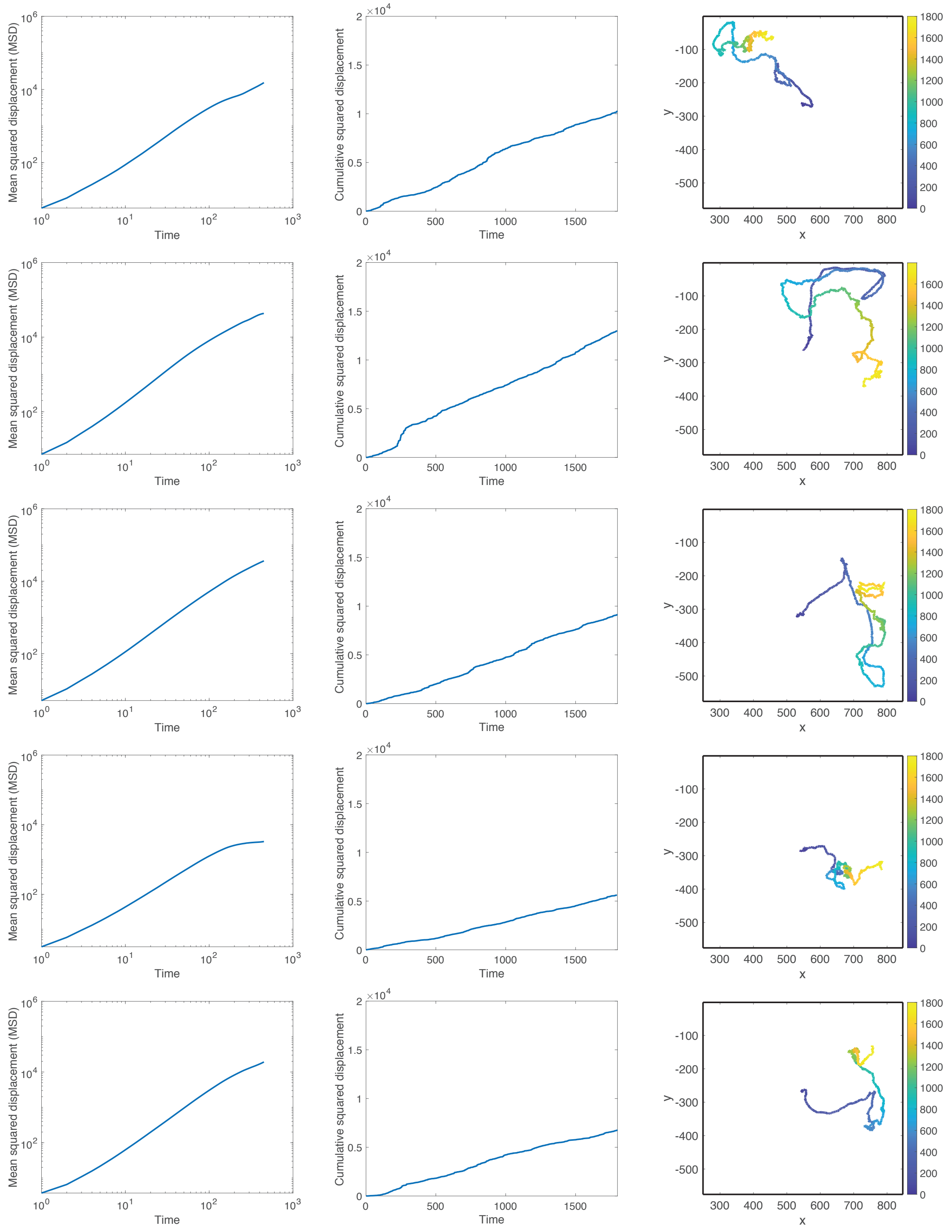


Figure 8: **Single worm experiments at $T = 26°C$.** Left column: mean squared displacement; middle column: cumulative squared displacement; right column: trajectory of worm. Each row represents a different 15-minute trial.

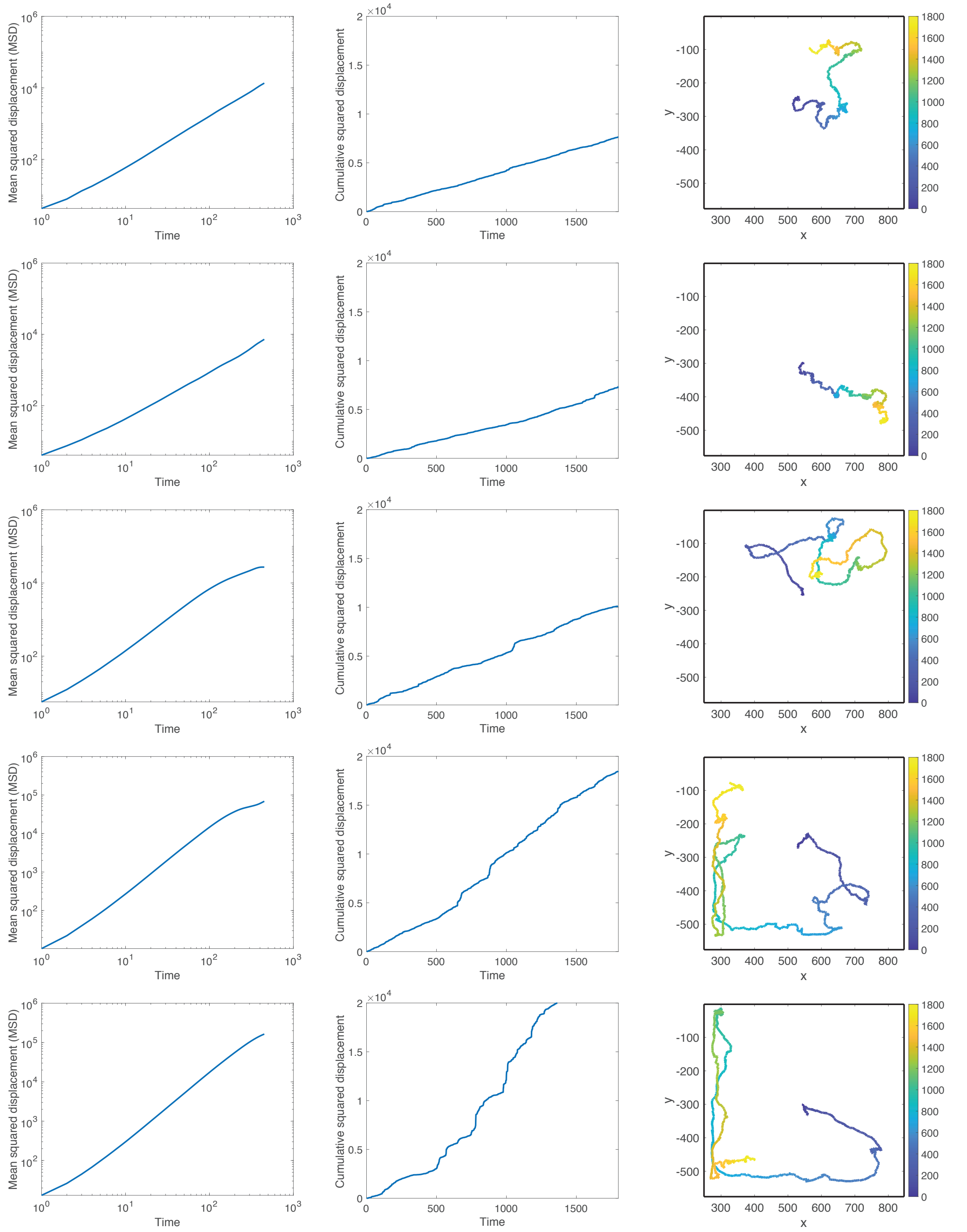


Figure 9: **Single worm experiments at $T = 28$°C.** Left column: mean squared displacement; middle column: cumulative squared displacement; right column: trajectory of worm. Each row represents a different 15-minute trial.

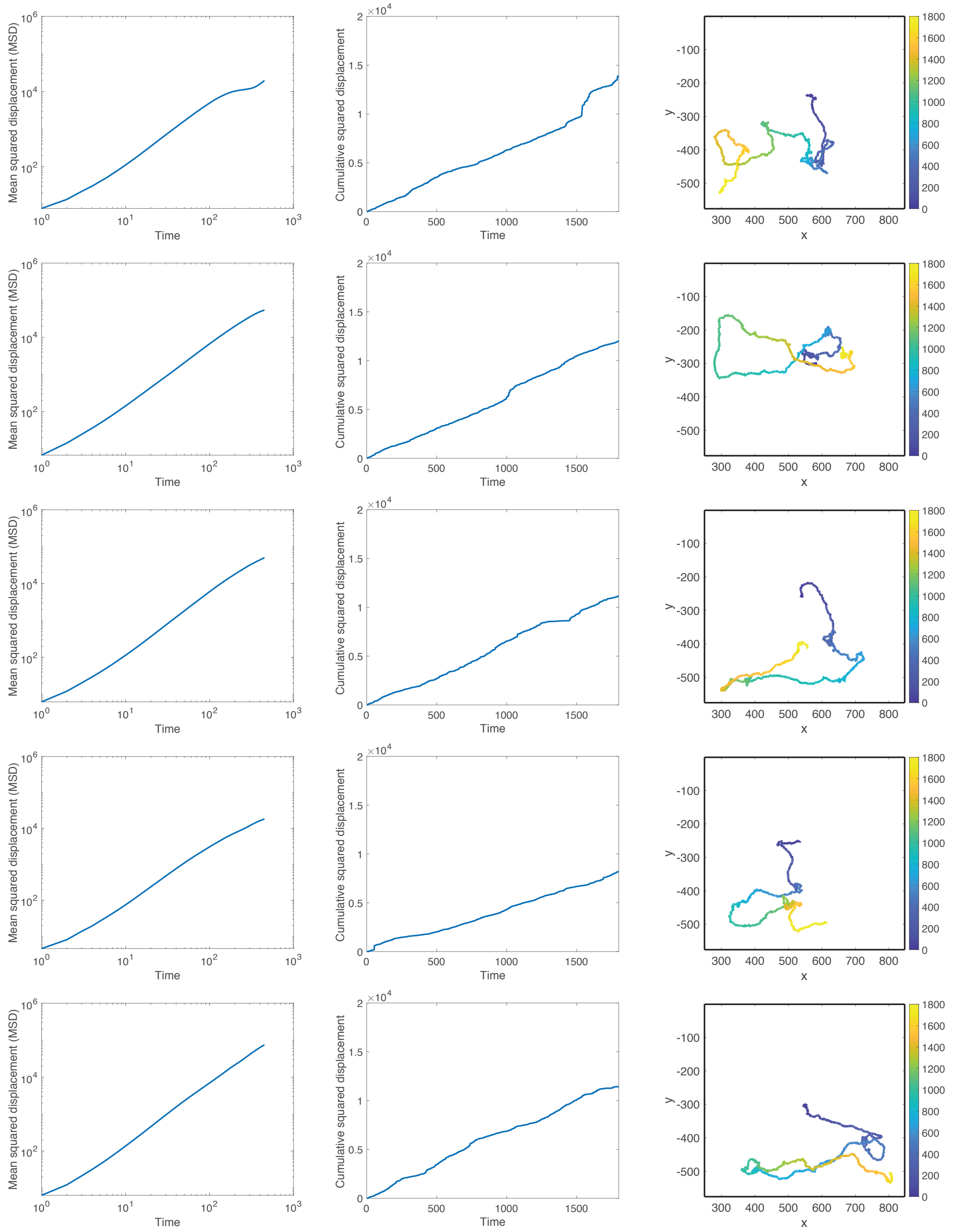


Figure 10: **Single worm experiments at $T = 30°C$.** Left column: mean squared displacement; middle column: cumulative squared displacement; right column: trajectory of worm. Each row represents a different 15-minute trial.

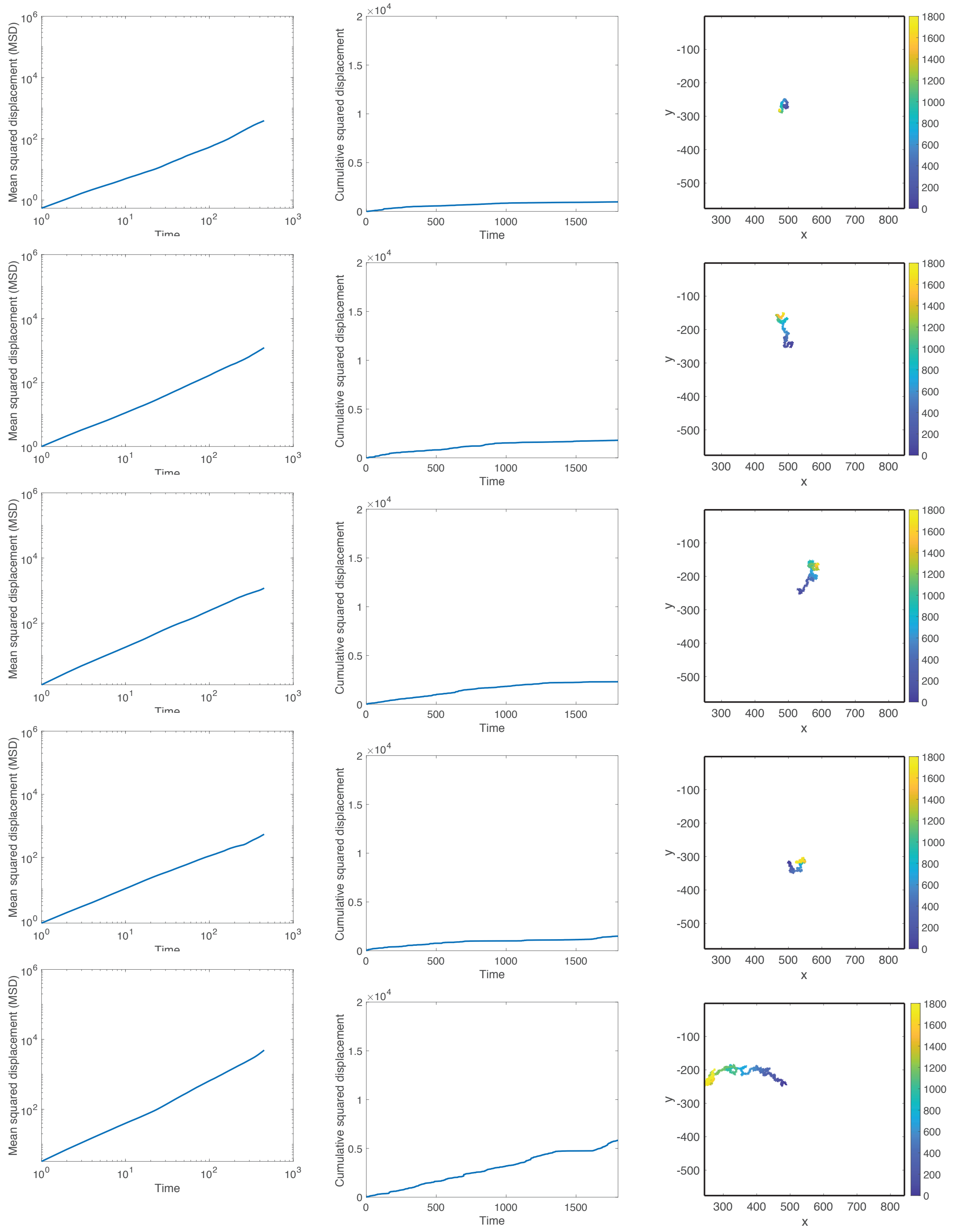


Figure 11: **Single worm experiments at $T = 32$°C.** Left column: mean squared displacement; middle column: cumulative squared displacement; right column: trajectory of worm. Each row represents a different 15-minute trial.

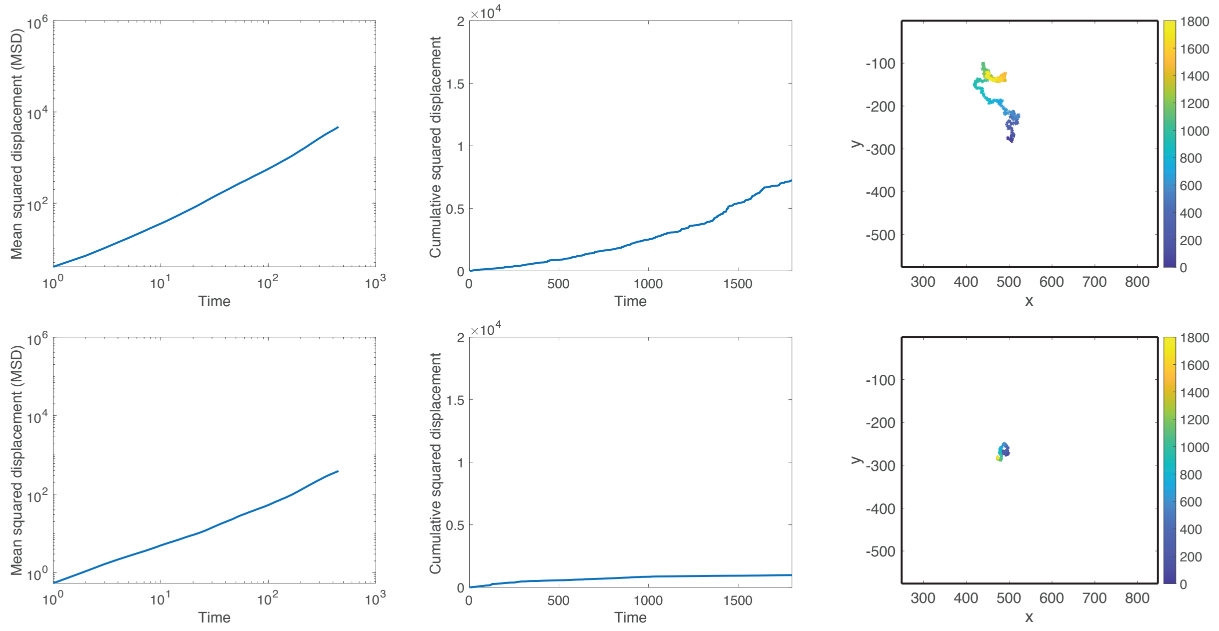


Figure 12: **Single worm experiments at $T = 34°$C.** Left column: mean squared displacement; middle column: cumulative squared displacement; right column: trajectory of worm. Each row represents a different 15-minute trial. Only two trials were performed due to the high temperature being detrimental to the health of the worms.
